# Strengthening Global Trade Regulation Through Targeted Listings on CITES Appendix III

**DOI:** 10.1101/2025.05.06.652200

**Authors:** Freyja Watters, Phillip Cassey

## Abstract

Appendix III is an underutilized component of The Convention on International Trade in Endangered Species of Wild Fauna and Flora (CITES), comprising less than 2% (n = 519) of all listed species (January 2024). Appendix III uniquely offers unilateral protection for nationally protected species without requiring international consensus.

Since CITES came into effect in 1975, 2,203 species have been added to Appendix III, including 875 endemic and 1,328 multi-country species. A total of 994 species have been delisted, mostly during CITES’ first decade. One-third of Appendix III species have been proposed for uplisting, with a 96% success rate and a median time of 3.3 years. Strengthening Appendix III’s impact involves broader cooperation for species ranging from multiple countries, regular reviews, and utilizing Appendix III listings as a precursor for Appendix I and II proposals to enhance global biodiversity management.

## Introduction

The Convention on International Trade in Endangered Species of Wild Fauna and Flora (CITES) is an international agreement that aims to ensure international wildlife trade does not threaten species survival. CITES governs over 40,000 species through three Appendices, each providing different levels of regulation. Appendix I enforces the strictest controls, prohibiting commercial trade unless from an approved captive breeding or artificial propagation program. Whereas, Appendix II species can be traded commercially, whether sourced from the wild or captivity. Trade in both Appendices I and II species requires an export permit or re-export certificate, and an additional import permit for Appendix I species. Export permits should be issued only after two evaluations by national CITES authorities: a Legal Acquisition Finding, and a Non-Detriment Finding (NDF). These assessments help ensure that trade is legal and does not jeopardize species’ survival. All decisions on Appendices I and II species are made at the regular Conference of the Parties (CoP) meetings, requiring a two-thirds majority by present and voting member states (known as Parties) for approval.

Appendix III has specific trade and listing requirements that differ from those of the other two Appendices. It is designed to support species that are protected within a Party’s borders, where that Party seeks international cooperation to help regulate trade. The inclusion or exclusion of species in Appendix III is at the discretion of the listing Party, offering greater autonomy than the collaborative decision-making process required for Appendices I and II. Trade regulations for Appendix III are also less stringent and do not require NDFs. For exports, the listing Party must issue a CITES export permit, confirming the specimen was not taken in contravention of national laws. For trade from non-listing Parties, CITES documentation requirements vary (see Table 1).

**Table 1:**
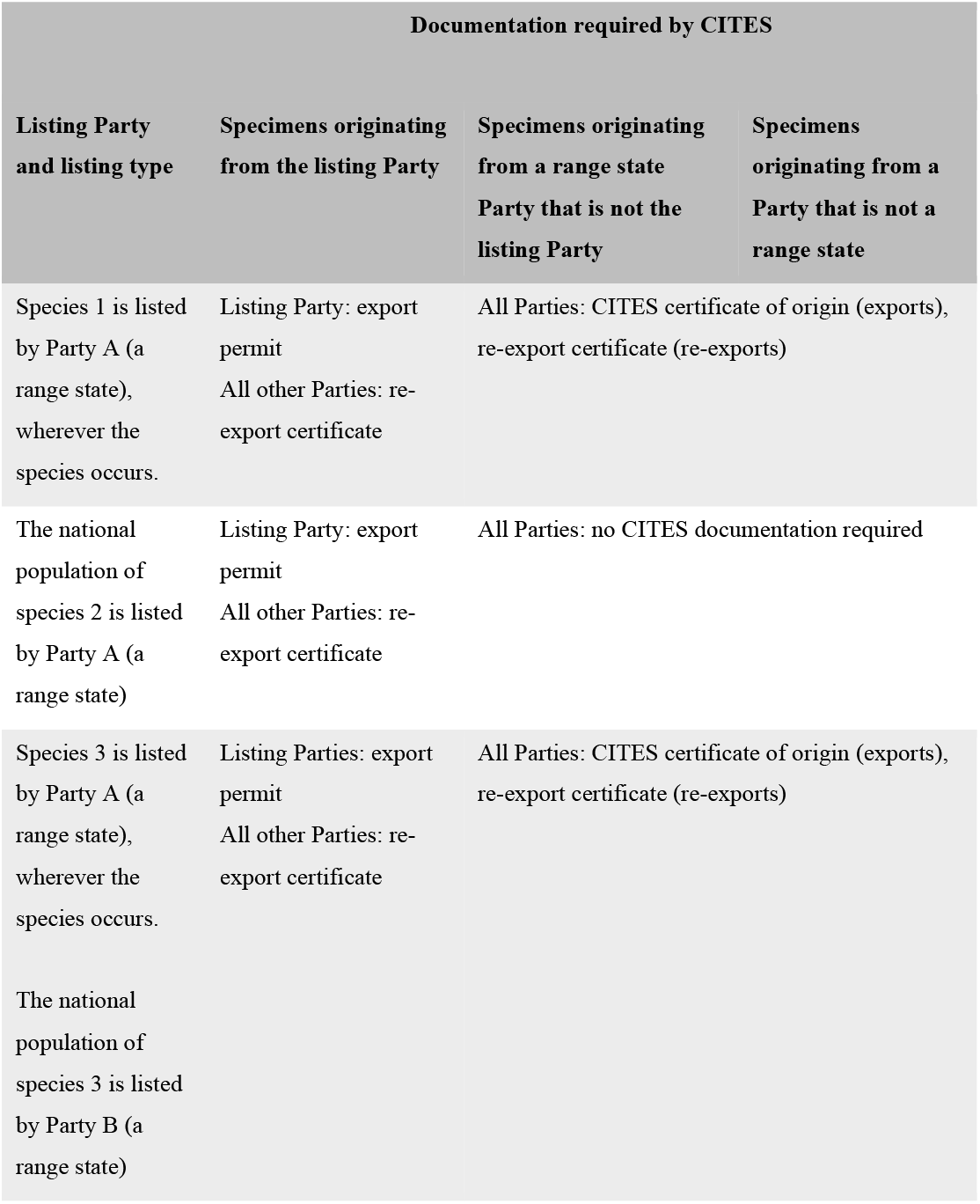
CITES trade documentation requirements for species listed in Appendix III.

CITES Appendix III comprises only 1.3% (n = 519) of all listed species (current January 2024). Resolution Conf. 9.25 (Rev. CoP17 and Rev. 18) highlighted several concerns, which may explain the low percentage of species listed in Appendix III, namely that: i) Appendix III currently contains species that rarely or never occur in international trade; ii) Parties are unwilling to take on the administrative burden of implementing the provisions of Appendix III; and iii) this unsatisfactory implementation arises because the Parties are not fully convinced of the effectiveness of Appendix III. In addition, for species with ranges spanning multiple Parties’ jurisdictions, traders often lack clarity on which populations are protected by Appendix III and what documentation is needed (see CoP15 Doc. 67).

Including species in Appendix III can strengthen domestic regulations and provide a framework for international trade cooperation (Heinrich et al 2022a). Member Parties are required to adopt national legislation to implement the convention, including laws that prohibit imports of species listed in any of the three Appendices without the correct documentation (Resolution Conf. 8.4 (Rev. CoP15)). Without CITES, national laws generally only protect native species, meaning that many Parties lack the legal framework to refuse imports of non-native species not listed in CITES (UNODC, 2020). Appendix III listings can strengthen domestic regulations as an immediate response for threatened species, without the need for international consensus, as is required for Appendix I and II listings.

CITES Parties have called for a more effective and consistent implementation of Appendix III (see CoP18 Doc. 100). To address this need, we explored the historical use of Appendix III, identified its most effective applications, and suggested ways to enhance its future effectiveness. Given concerns about non-traded species in Appendix III, we analyzed the frequency of species listings and delistings, focussing on whether deletions involved non-traded species. Our analysis included a review of endemic and non-endemic species listings and investigated how Appendix III has curbed illegal trade by examining seizure data over time.

We investigated the success rate of species proposed for uplisting to evaluate how effectively Appendix III supports proposals to the other two Appendices.

CITES lacks a formal review for Appendix III listings, essential for effective trade regulation. We analyzed current Appendix III listings and trade data to identify species either not involved in trade or facing high trade volumes, indicating potential unsustainable trade threats. This information is critical for setting conservation priorities and determining if a species should remain in Appendix III, be delisted, or be considered for uplisting for greater protection.

## Methods

Using the “ History of Listings” dataset from the Checklist of CITES Species website (checklist.cites.org), we compiled historical listings for Appendix III up to January 2024. We used the “ Full species list” on the Checklist website to gather data on range countries for each species and obtained International Union for Conservation of Nature (IUCN) Red List status from iucnredlist.org (IUCN 2023). Trade volumes and seizures were estimated by extracting shipment-level export data (excluding re-exports) from the CITES Trade Database (trade.cites.org).

Data on Appendix I and II proposals were compiled from the CITES CoP meeting web portal (cites.org/eng/meetings/cop). Manual searches of relevant web pages and PDFs provided specific data on proposed species, Appendices for amendment, and final decisions. We used a modified Stage 1a of the CITES Review of Significant Trade (RST) process to identify noteworthy trade patterns in Appendix III species. We extracted annual report statistics for the five most recent, complete years of data (2018-2022) from the CITES Trade Database focusing on direct exports of wild, ranched, or unknown origin. The analysis followed the RST criteria in Resolution Conf. 12.8 (Rev. CoP18) with adjustments for Appendix III species. Further methodology outlining the modified RST data and selection criteria used can be found in Table S1.

Statistical analyses were conducted using the R software environment for statistical and graphical computing (ver. 4.4.0) (R Core Team, 2023). We analyzed the proportion of endemic, and non-endemic species listed in Appendix III using a beta-binomial GLMM approach with the glmmTMB function in the glmmTMB package to assess the proportional distribution changes over time with count data (Brooks et al., 2017). Trends in seized shipments of Appendix III species were examined using a negative binomial GLM with the glm.nb function from the MASS package (ver. 7.3-60.0.1) (Venables & Ripley, 2002). Post-hoc tests were conducted using the emmeans package (ver. 1.10.0) (Lenth, 2024). Time-to-event analysis for uplistings from Appendix III to I or II was conducted using the Kaplan-Meier method and log-rank test, with analyses performed using the survfit and survdiff functions in the survival package and plotted with the ggsurvfit function (ver. 3.5.7) from the ggsurvfit package (Sjoberg et al., 2023; Therneau, 2023).

## Results

Since CITES came into effect in 1975, 2,203 unique species have been added to Appendix III (Fig. 1), including 875 endemic and 1,328 multi-country species. Only four multi-country species were listed by all native range Parties (Table S2-S4). 994 species have been delisted, mostly from listings in CITES’ first decade (Fig 2, Fig S1). Of the early deletions, 804 were species listed by Ghana in 1976, which were delisted in 1985 following Resolution Conf. 5.22, which mandated that species must be native to the listing Party and protected under their national legislation. Before this delisting, an average of 3.4% of listed Appendix III species were found in trade annually, but this rose to an average of 44.4% between 1986-2019 (Fig. 2). Ghana delisted all remaining Appendix III listings in 2007, 88% of which were actively involved in trade, including many heavily traded species (Table S5). This included 34 of the top 50 wild-caught live CITES bird species traded between 2003-2007 (Table S6).

**Figure 1.**
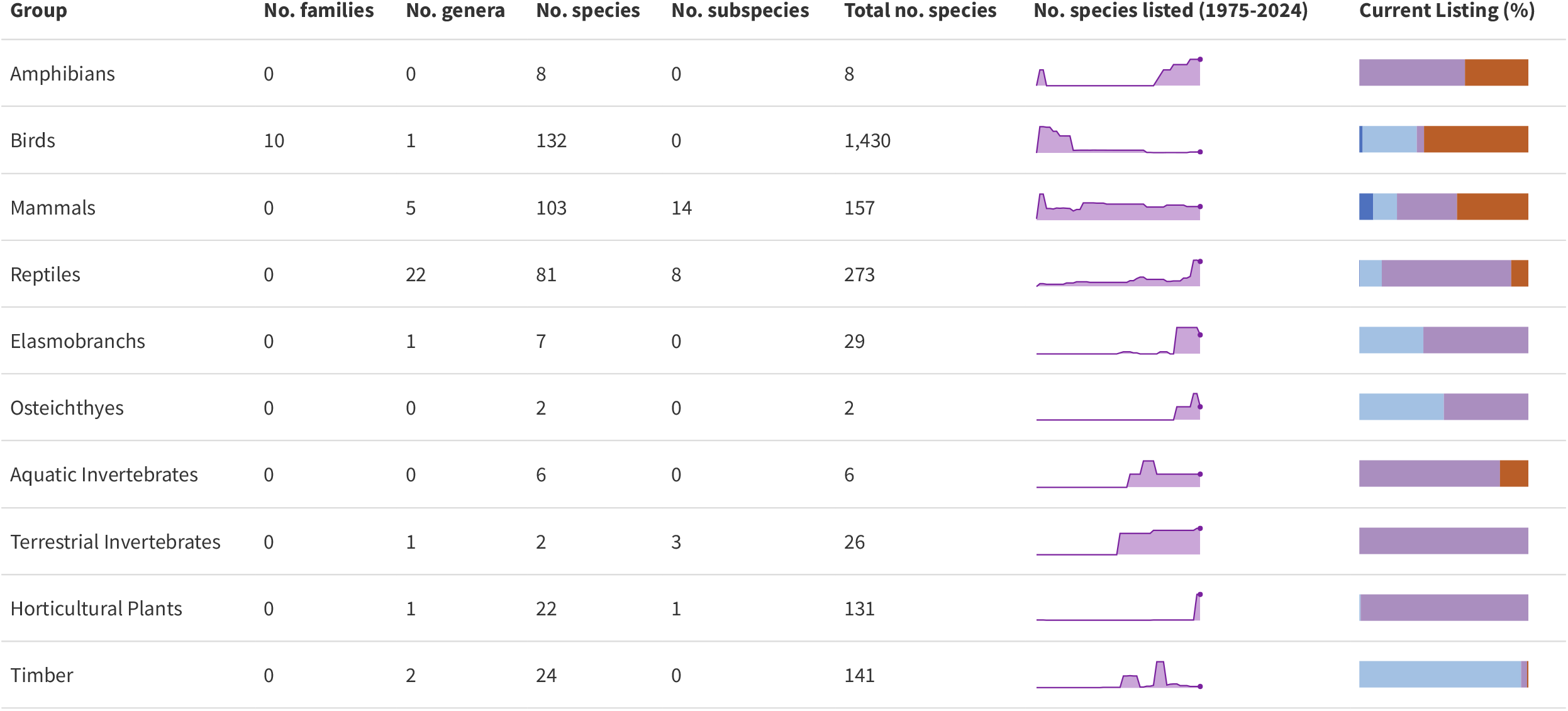
The total number of species ever listed in Appendix III. The area chart shows the number of species listed in Appendix III through time. Horizontal coloured bars represent the listing status of species as of January 2024, expressed as percentages of the total number of species initially listed in Appendix III. Dark blue represents species listed in Appendix I, light blue for Appendix II, purple for current Appendix III listings and red for species removed from all CITES Appendices

**Figure 2.**
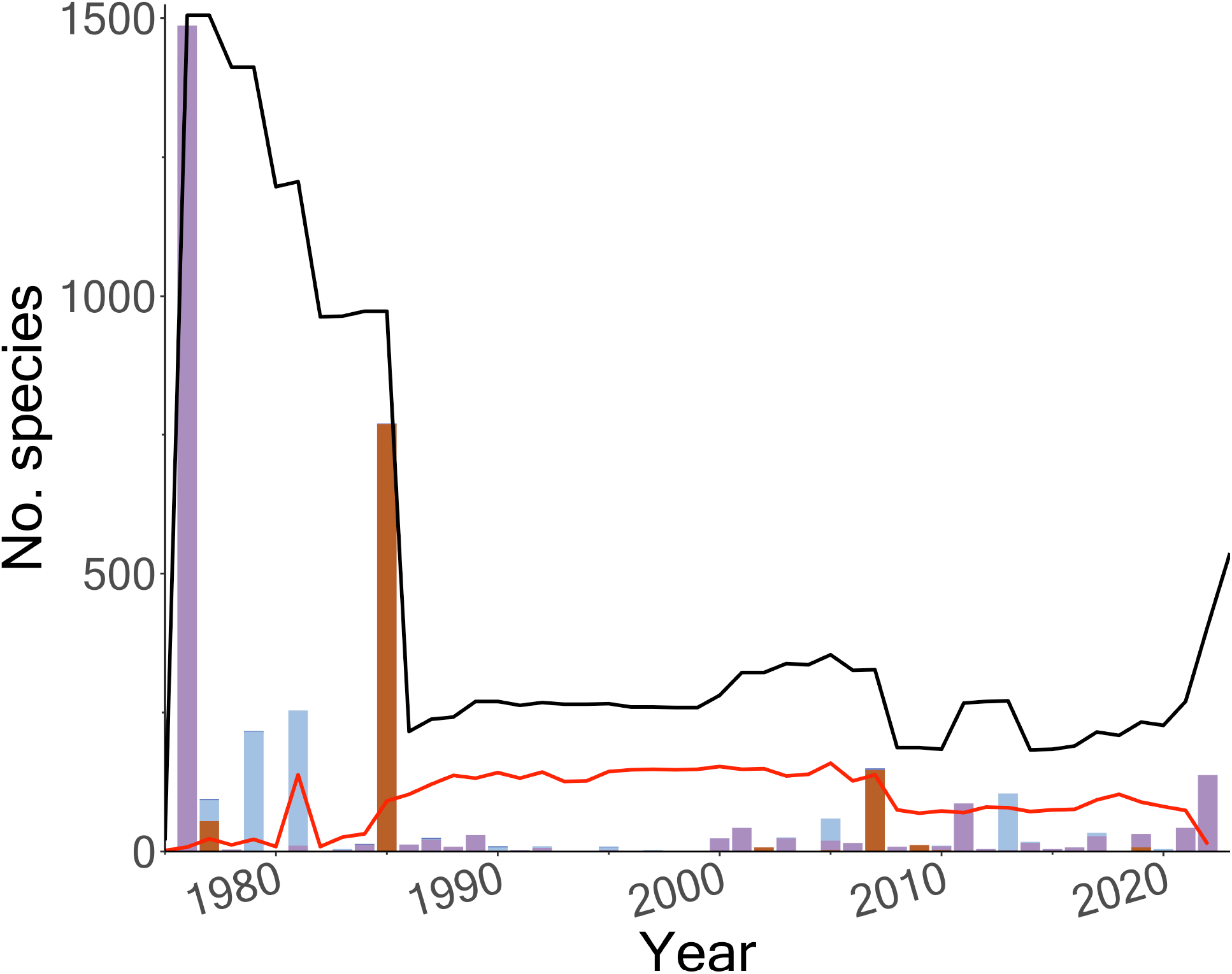
Total number of species listed in Appendix III through time (black line), the number of listed species present in CITES trade each year (red line) with coloured columns showing new Appendix III listings (purple), uplistings from Appendix III to Appendix I (dark blue) and II (light blue) and the removal of Appendix III species (red) from CITES by listing Parties.

Over time, there has been a significant increase in Appendix III endemic species listings and a decrease in non-endemic listings (Fig. 3). As of January 2024, 68% of Appendix III species are endemics.

**Figure 3.**
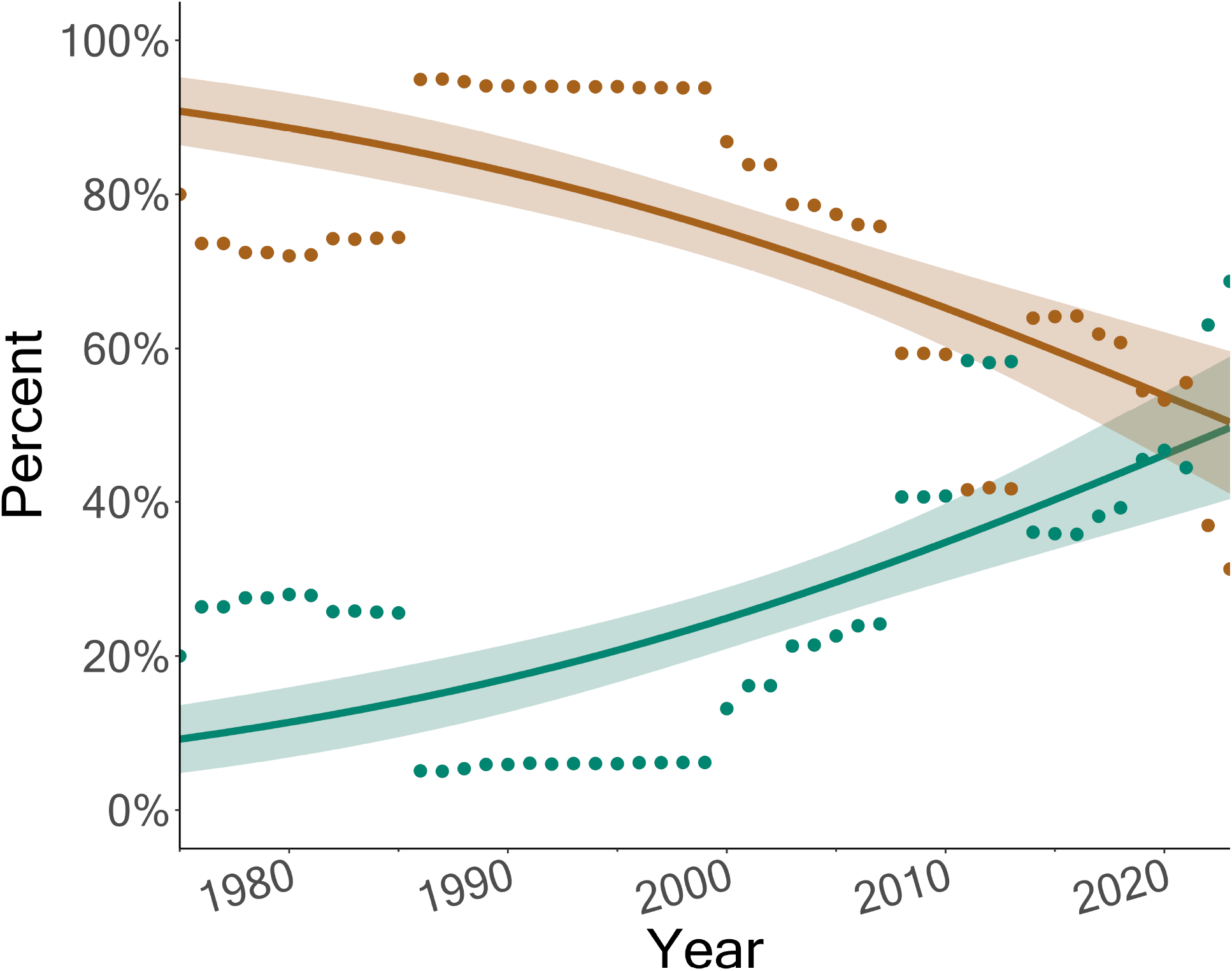
The percentage of Appendix III species that were endemics (dark green), and non-endemics (brown). Points show actual values, while solid lines and shaded areas represent the mean predicted values and 95% confidence intervals for the proportion of listings over time, as derived from beta-binomial generalised linear mixed models (GLM-NB) which showed a significant increase in endemic species listings alongside a significant decrease in multi-country species listings over time (df = 1, χ^2^ = 62.148, p < 0.001***)

Seizures of Appendix III shipments increased significantly, peaking in 2019, (Fig. 4). Pairwise comparisons indicated that listing Parties experienced significantly fewer seized shipments compared to exports originating from both non-range Parties, and non-listing range Parties (Fig. 4).

**Figure 4.**
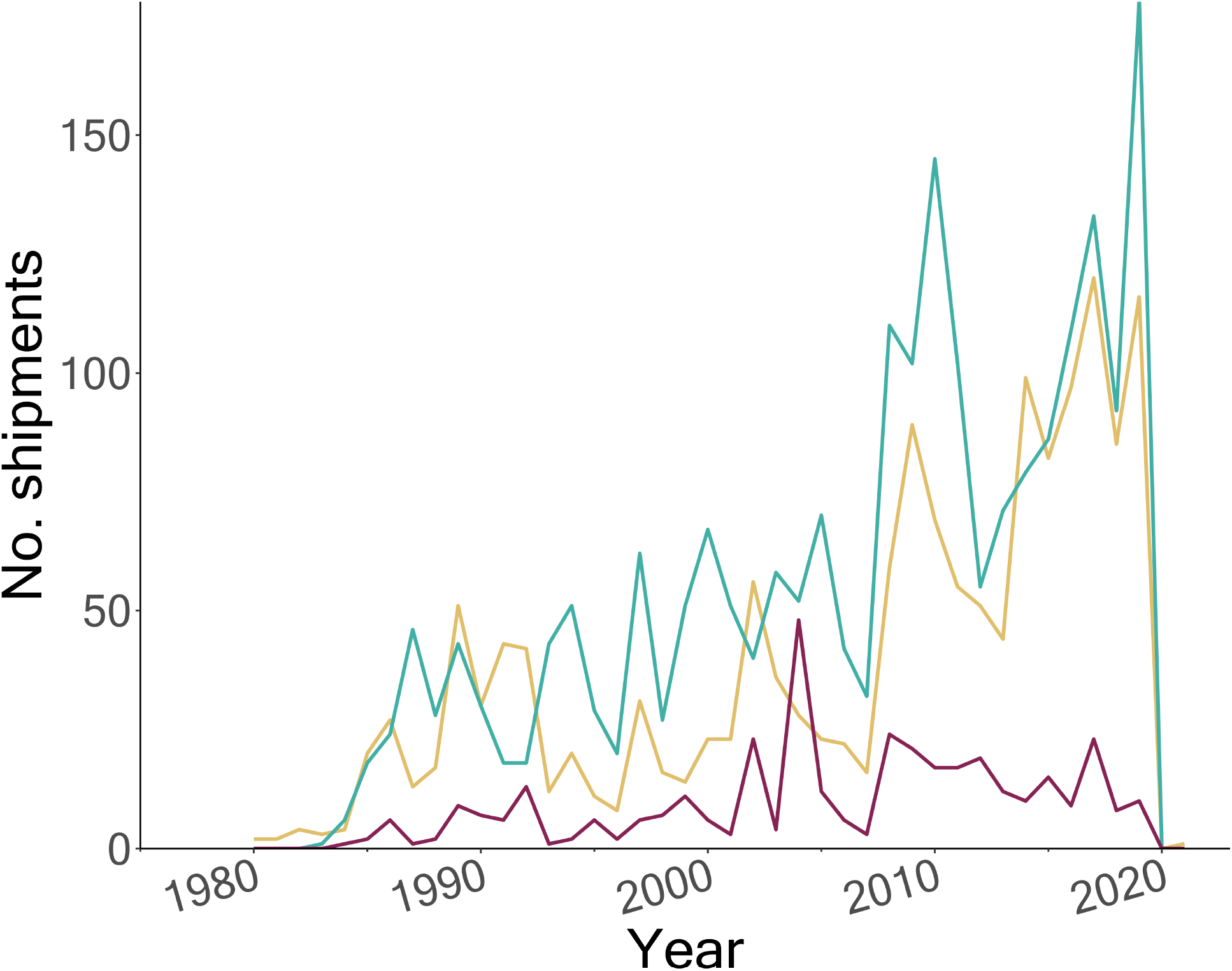
The annual count of seized shipments containing Appendix III species, as recorded in the CITES Trade Database, originating from countries categorized as either: a) the listing Party for the shipped specimens (dark red); b) a native range Party but not the listing Party for the shipped specimens (yellow); or c) neither a native range nor a listing Party for the shipped specimens (teal). Beta-binomial generalized linear mixed models (GLM-NB) showed that seizures of Appendix III species shipments increased significantly, peaking in 2019, before declining (estimate = 0.1, SE = 0.01, z-value = 20.9, p < 0.001). Post hoc pairwise comparisons on the GLM-NB model indicated that listing Parties experienced significantly fewer seized shipments compared to exports from both non-range Parties (estimate = 1.4, SE = 0.2, z-ratio = 7.1, p < 0.001) and range nations that are not listing Parties (estimate = 1.7, SE = 0.2, z-ratio = 8.8, p < 0.001). There was no significant difference in seizure numbers between non-range Parties and range nations that are not listing Parties (estimate = -0.3, SE = 0.2, z-ratio = -1.7, p = 0.2).

One-third (n = 730) of Appendix III species have been proposed for uplisting to Appendix I or II, with 96% successfully uplisted (Fig S2). Specifically, 37 species moved to Appendix I, 5 were split-listed (I/II), and 657 were uplisted to Appendix II. Most species (n = 675) were successfully uplisted on their first attempt (Fig S2). Several species were added to Appendix III after a failed Appendix I or II proposal, with the median lag time between an unsuccessful proposal and Appendix III listing being 1.5 years (Fig. S3). Of the species added to Appendix III following an unsuccessful proposal, 24 out of 42 were later uplisted to Appendix II at subsequent CoP meetings. For species successfully uplisted, the median time from Appendix III listing to uplisting was 3.3 years (Fig 5).

**Figure 5.**
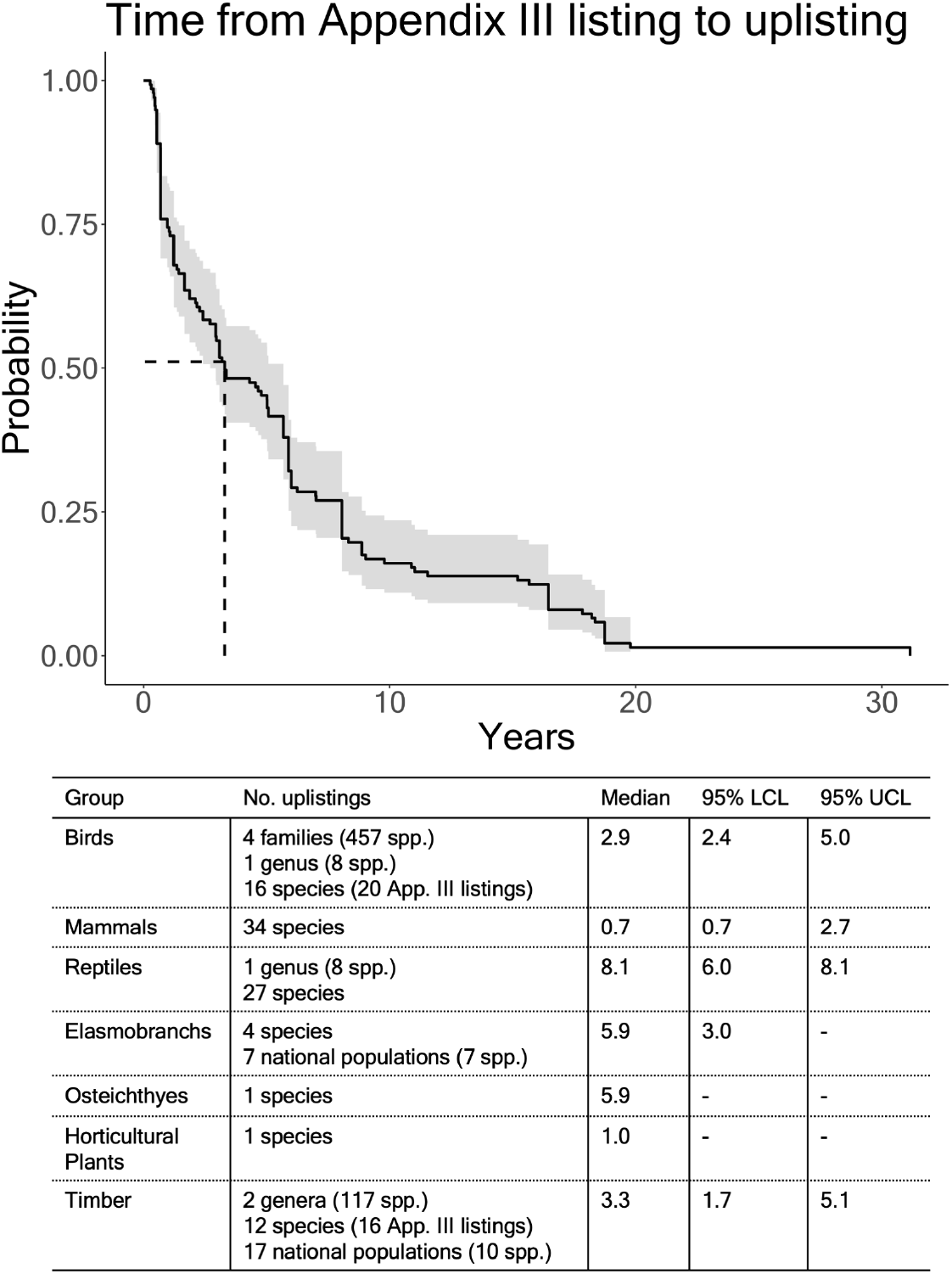
Kaplan-Meier plot showing the lagtime in years between an Appendix III listing and the subsequent uplisting of the species to Appendix I or II at a Conference of Parties (CoP) meeting (for all successfully uplisted species). The shaded band represents 95% confidence intervals. The median time between events is shown as a dashed line (3.3 years). The median time between uplistings did not significantly differ across taxonomic groups (log-rank test, χ2 = 5.4, df= 6, p = 0.4), Species uplisted together in a single higher taxonomy proposal (e.g., a genus or family listing) are considered as one data point.

Appendix III lists 519 species (January 2024), (Table S2). Among these, 88 species (17%) have been listed for over 30 years, including 13 species listed since 1975, whereas 312 species (60%) were added within two years of the most recent CoP (CoP19, 2022), (Table S2). These recent additions have not yet entered trade (Table S2). Of the other 205 species, 97 have been involved in trade in the last five years (2018-2022), 5 genera (28 spp.) and 27 species have both been listed for over a decade and not traded in the previous decade (2012-2022) (Table S2). Twenty species found in trade in the past five years exhibited significant trade patterns as identified by the RST selection criteria (Table S7). Among these, eight species were classified as threatened or near-threatened by the IUCN, with five fulfilling the criteria for High-volume (globally threatened) selection.

## Discussion

In CITES’ first decade, Appendix III was overburdened with non-traded species, an issue largely resolved by Ghana’s 1985 delisting. Presently, the concern that Appendix III is ineffective due to the listing of species not in international trade (e.g. Resolution Conf. 9.25 (Rev. CoP17 and Rev. 18) is largely unjustified. An average of 44.4% of Appendix III species were traded annually until 2019, which is noteworthy considering that only 30% of Appendix II species were traded between 2011-2020 (CITES Secretariat, 2022). The further reduction in traded species richness between 2020-2022 was likely due to delays in annual reports and the COVID-19 pandemic (CITES Secretariat, 2022). Listed species not frequently appearing in trade are often endemic and were listed to curb illegal trade, with legal export prohibited, or are part of higher taxonomy listings to avoid look-alike issues and misidentification. However, we recommend that for listings active for a decade without trade or risk, the listing Party should consider evaluating what benefits a continued listing offers.

The increase in endemic species listings reflects a growing recognition of the benefits of Appendix III listing, especially in combating illegal trade. Endemic species, unlike those found in multiple countries, avoid illegal trade through origin misrepresentation; because they originate from a specific location, it is harder for traders to misrepresent them as being from another country. This clarity simplifies the implementation of trade regulations. For Parties with nationally protected endemic species, facing threats from illegal cross-border trade, Appendix III can reinforce domestic conservation laws by ensuring importing Parties are equipped to identify and seize non-compliant shipments (Heinrich et al., 2021; Heinrich et al., 2022b; Wijnstekers, 2018).

Listing non-endemic species in Appendix III presents challenges due to their distribution across multiple jurisdictions and the varying degrees of threats they may face. Historically, collaboration between range Parties has been minimal, with less than 1% of Appendix III species native to multiple countries listed by all relevant Parties. Ghana’s 2007 delisting highlights the limitations of single-Party listings for non-endemics. Their delisting included 73 songbird species, which accounted for 70.7% of all CITES live wild bird trade before removal, and led to a perceived decline in global trade initially attributed to the EU’s wild bird trade ban (Juergens et al., 2023). This finding underscores the importance of Appendix III listings for data collection and the impact of unilateral delisting, as without CITES regulation, there is no international species-level trade tracking, obscuring trade trends. This may increase the risk of overexploitation and illegal trade in vulnerable species. Parties considering listings for multi-country ranging species should involve discussions with other range Parties.

Increased seizures from non-listing Parties suggest that Appendix III can be effective in curbing illegal activities. Conversely, fewer seizures in trade originating from listing Parties indicates that Appendix III enhances national efforts to prevent illegal cross-border trade. Non-listing range Parties should consider Appendix III to access international regulatory tools and support, enhancing their capacity to detect and deter illegal trade.

### From Initial Listing to Uplisting: A Pathway to Stronger Protection

Since 1975, 96% of species proposed for uplisting from Appendix III to either I or II have been successful. Appendix III is useful when trade information is initially missing, allowing for subsequent data collection to support a higher listing. For example, the United States preemptively listed species from the Apalone genus in Appendix III in 2016, anticipating increased demand based on the “ boom-and-bust” cycles in turtle harvesting (CITES, 2022). Subsequent trade increases in Apalone spp. supported their uplisting to Appendix II in 2022, to mitigate overexploitation.

Appendix III listings have served as a secondary measure after failed Appendix I or II proposals. For several commercial timber species like *Cedrela odorata, Dalbergia retusa, D. stevensonii*, and *Swietenia macrophylla*, interim Appendix III listings have bridged the gap between Appendix II uplisting attempts. Challenges and misconceptions about CITES listings for timber species have historically hindered Appendix II support (Reeve, 2015). As an interim measure, Appendix III bolsters international support by enhancing national regulations, showing Party commitment, and generating supporting trade data (Buitrón & Mulliken, 2003; CITES, 2013).

An Appendix III listing allows for immediate action, which can be crucial when waiting 2-3 years for an unguaranteed Appendices I or II inclusion would leave Parties without means to regulate cross-border trade. Parties should consider this proactive approach for species facing trade risks. The earless monitor *Lanthanotus borneensis* was proposed for Appendix I in 2016 (but amended to Appendix II with a zero wild quota) the species faced increased smuggling and market value due to the delay in protection between the proposal announcement and CoP decision (Janssen & Krishnasamy, 2018). The median time between a species Appendix III listing and its uplisting to Appendix I or II was roughly equal to the interval between CoP meetings, suggesting that Parties can use this strategy to increase their chances of uplisting. However, it should be used more frequently. At CoP19, only one of 28 proposals for new CITES listings used an interim Appendix III listing (Australia’s Appendix I submission for *Tiliqua adelaidensis*). Parties planning future proposals could benefit from this approach to deter a rise in illicit trade demand during the interim period triggered by CoP proposals.

### Review mechanism for Appendix III listings

There is no formal trade review mechanism for Appendix III, with the responsibility of periodical review falling to the listing Party. However, many national CITES authorities face funding shortages and may lack the resources to consistently review trade data (Wyatt, 2021). Standardizing the review of Appendix III species could be achieved by incorporating it into the existing Review of Significant Trade (RST) process. Since Appendix III listings are voluntary, including a review of Appendix III trade in the initial trade analysis (Stage 1a) of the RST would alert Parties to significant Appendix III trade volumes, and potentially unsustainable trade activities and signal to other range Parties the need for increased national protections or their own listing.

To demonstrate this approach, we applied the extended Stage 1a analysis of the CITES RST process to Appendix III species, investigating significant species/export-country trade patterns between 2018-2022. The highest traded Appendix III species was the brown sea cucumber, *Isostichopus fuscus*, which is Red Listed as Endangered. Native to 10 Parties’ coastal waters, it is listed in Appendix III only by Ecuador, where overfishing has nearly depleted it (Conand, 2017). CITES records from 2022 showed Ecuador exported 547,039 whole cucumbers, and 90,734 under the ambiguous “ specimen” code, typically reserved for scientific samples, but likely representing whole cucumbers given their commercial purpose. Although CITES export data may contain inaccuracies (Berec et al., 2018), the fact that Ecuador’s exports were at or slightly over the country’s annual quota of 600,000 underscores national authorities’ need for ongoing vigilance. Additionally, this Appendix III data provides transparency to national quota figures, highlighting the role of international regulation in assisting compliance with national measures. Pursuing an Appendix II listing, which this species appears to fulfil the criteria for, would greatly assist in promoting research into sustainable yields for NDFs, ensuring sustainable trade levels.

In the interim, supporting Appendix III listings by other range Parties, particularly major exporters like Mexico, who face similar challenges from overfishing and unsustainable catch practices, is essential (Herrero-Pérezrul & Chávez, 2024).

## Conclusions

Appendix III is effective but underutilized. It uniquely allows countries to unilaterally protect and monitor native species internationally without requiring consensus, a significant advantage frequently overlooked by many CITES Parties.

We recommend several steps to strengthen the impact of Appendix III listings. Firstly, broader cooperation is essential for species native to multiple Parties, ensuring comprehensive cross-border protection. For endemic species, Appendix III listings are beneficial when national laws are robust, with CITES listings empowering other Parties to confiscate illegal shipments effectively.

Appendix III listings provide critical monitoring data instrumental in building support for potential uplistings. Parties proposing Appendix I or II listings should additionally consider implementing an Appendix III listing before announcing proposals. This strategy acts as a safety net if the proposal fails and helps mitigate any increase in trade before the CoP decision.

To ensure listings effectively regulate trade, they must undergo regular review. Including Appendix III species in the RST process could facilitate this. This ongoing evaluation will help adapt and refine strategies to meet the changing dynamics of wildlife trade and conservation needs. Through these efforts, Appendix III can play a pivotal role in the sustainable management of global biodiversity.

## Acknowledgments

We acknowledge that the land on which we conducted our research is the traditional land of the Kaurna Meyunna of the Adelaide Plains. We pay our respects to Kaurna elders past, present, and emerging. This research was partly supported by an Australian Research Council Discovery grant (DP210103050) and Industry Laureate Fellowship ‘Combatting Wildlife Crime and Preventing Environmental Harm’ (IL230100175) to P.C. F.W. was supported by an Adelaide University Postgraduate Research scholarship and stipend.

