## Supplemental Figure 1 for "Strengthening Global Trade Regulation Through Targeted Listings on CITES Appendix III"

**Supplementary Figures**

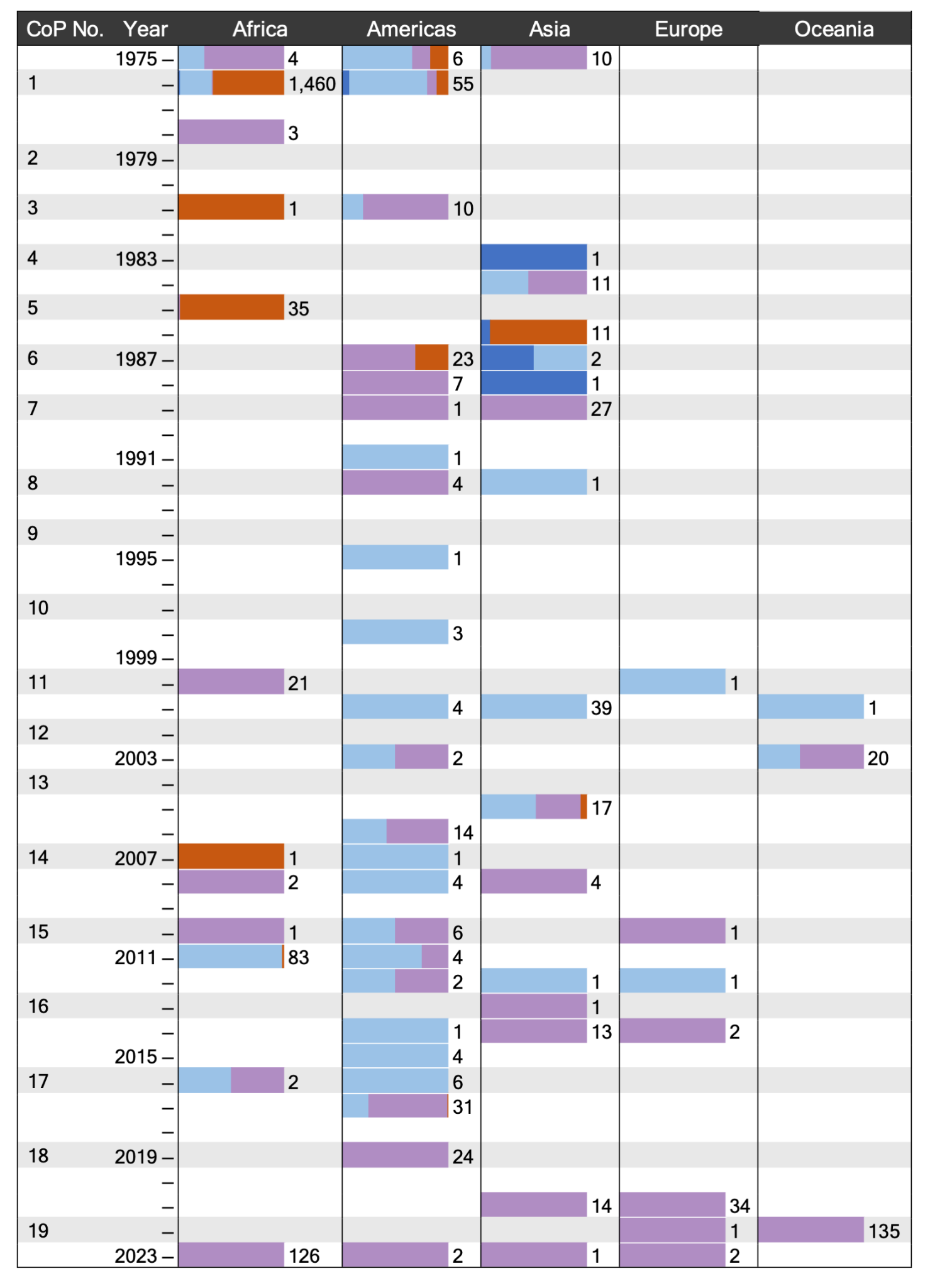

Figure S1. Species listed in Appendix III of the Convention on International Trade in Endangered Species of Wild Fauna and Flora (CITES), grouped by their listing date (year) and the geographic region of the listing Party. Note that one species can be listed by more than one Party. Grey rows indicate that a Conference of the Parties (CoP) meeting was held that year. Horizontal coloured bars indicate the current listing status (as of January 2024) of species, shown as a percentage of the total number of species initially listed in Appendix III for a given year/region, which is displayed numerically to the right of each bar.

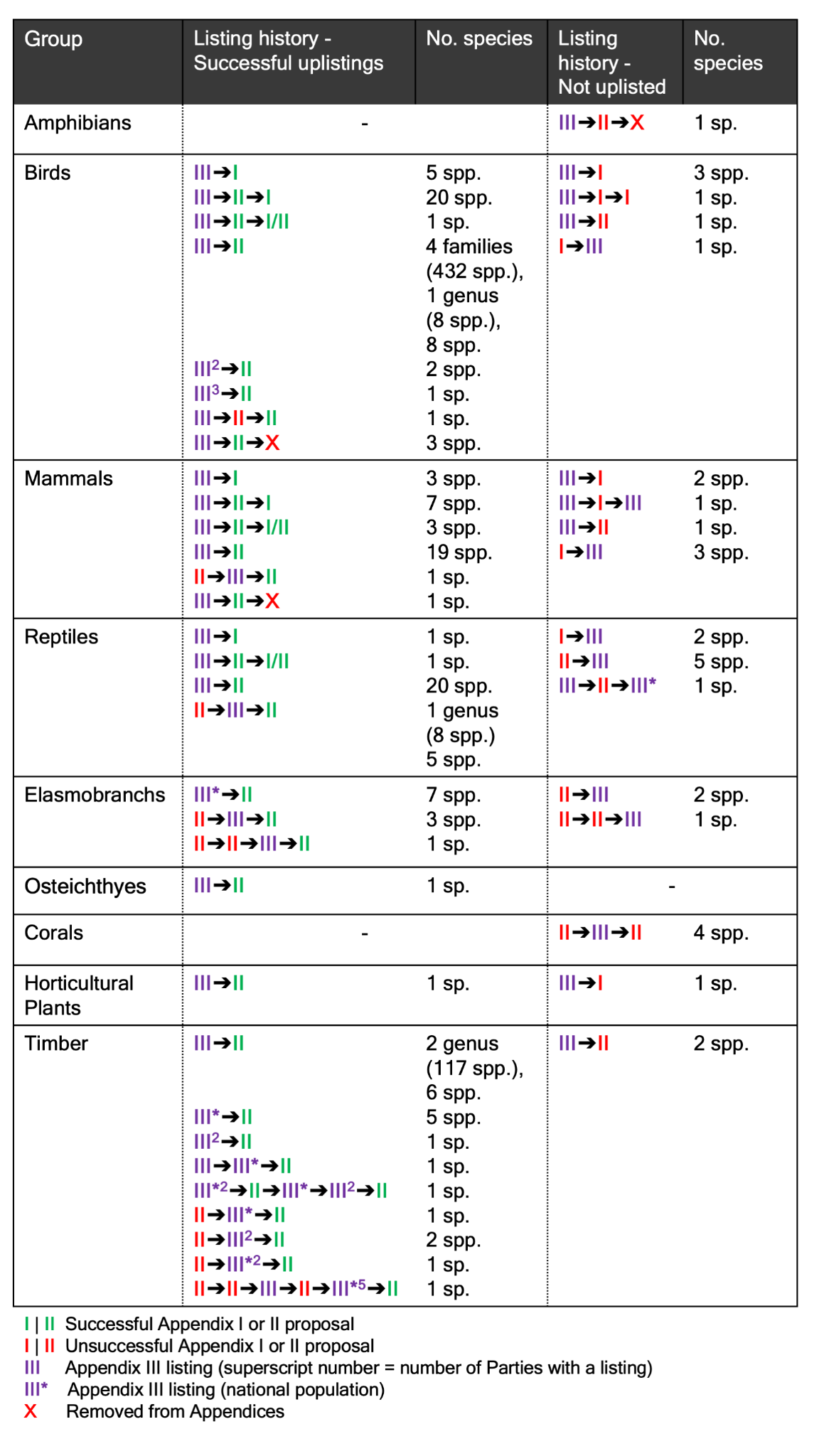

Figure S2. Species listed in Appendix III that have ever been attempted to be uplisted to Appendix I or II at a Conference of the Parties (CoP) meeting.

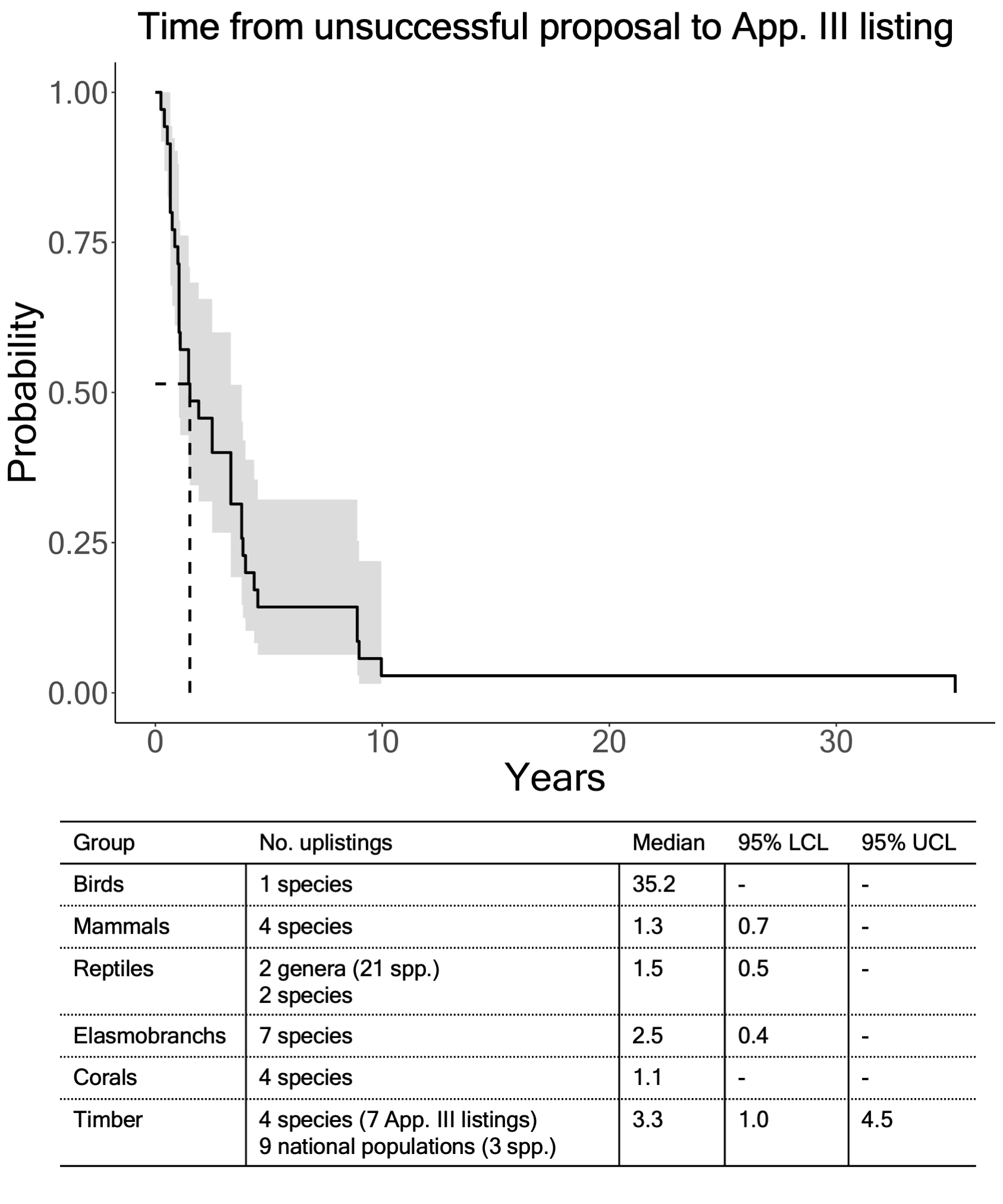

Figure S3. Kaplan-Meier plot showing lagtime in years between an unsuccessful Appendix I or II proposal at a Conference of Parties (CoP) meeting and a subsequent listing of the species in Appendix III. The shaded band represents 95% confidence intervals. The median time between events is shown as a dashed line. Species listed together as a single higher taxonomy listing (e.g. a genus or family listing) are counted as one data point.
